# Osteocytes secrete adiponectin and display adipocyte-like phenotype under control of PPARG nuclear receptor

**DOI:** 10.64898/2026.05.02.722443

**Authors:** Mohd Parvez Khan, Emily Crowe, Joshua Letson, Sudipta Baroi, Piotr J. Czernik, Beata Lecka-Czernik

## Abstract

Osteocytes and adipocytes represent cells with disparate functions. Osteocytes regulate bone metabolism (remodeling) and bone homeostasis, while adipocytes regulate energy metabolism and energy storage. Here, we demonstrate that osteocyte phenotype consists of adipocytic features which are under control of peroxisome proliferator-activated receptor gamma (PPARG), a master regulator of adipocyte differentiation and function. Using a mouse model with osteocyte-specific deletion of PPARG (OT^γKO^) and osteocyte cellular model of MLO-Y4 cells edited with CRISPR/Cas9 for PPARG deficiency, we are demonstrating that under PPARG control osteocytes produce and secrete adiponectin (ADIPOQ), and they are equipped in adipocyte-specific mechanisms for lipid-storage and their metabolism. Under PPARG, osteocytes accumulate lipid droplets which correlate with their capability to cover up to 20% of energy requirements from fatty acids metabolism. Although osteocytes like osteoblasts mainly express *perilipin 2* (*Plin2*), however similarly to adipocytes, lipid droplets accumulation is associated with expression of *perilipin 1* (*Plin1*) under PPARG control. Similarly, lipids accumulation and metabolism involve adipocyte-specific activities including *fatty acids binding protein 4* (*Fabp4*), *hormone-specific lipase* (*Hsl*) and *adipocyte-specific triglyceride lipase* (*Atgl*), which expression are under PPARG control. These studies provide a new understanding of osteocyte biology which include adipocyte-like endocrine and lipid metabolism features probably reflecting an adaptation to their unique localization and a need for a maintenance of functional fitness in these conditions. They deepen our comprehension of the crossroads of osteocyte and adipocyte function and underscore the therapeutic potential of targeting common molecular pathways in both cell types for managing metabolic disorders and skeletal diseases.

## INTRODUCTION

Osteocytes and adipocytes represent two distinct cell types of different function and location in the body. Osteocytes, the most abundant cells in bone, are multifunctional, dynamic, and through their paracrine/endocrine activities capable of communicating within the bone niche and with distant tissues. In bone, osteocytes control bone remodeling or metabolism, both directly via osteokines, or indirectly through regulation of marrow microenvironment, marrow adiposity, hematopoiesis, and response to biomechanical challenges ^1,2^. For a whole lifespan they reside in the lacunae of mineralized bone compartment with a connection to the environment limited to the system of canaliculi through which nutrients, oxygen, signaling and waste clearing are provided. In contrast, adipocytes are in direct contact and are highly responsive to the environmental cues since they function as specialized cells dedicated to energy storage and its dissipation. In response to energy demands, they either uptake triglycerides to store lipids in the process of lipogenesis or release free fatty acids in the process of lipolysis ^3^. Similarly to extramedullary adipocytes, bone marrow adipocytes also store lipids although this function is controlled by mechanisms specific to bone including their unique responses to WNT signaling, sympathetic nervous system (SNS) control, and nutritional cues^4,5^.

However, besides apparent differences in function, osteocytes and adipocytes share some similarities. Both originate from mesenchymal progenitor cells. They are non-proliferating terminally differentiated cells with a long lifespan counted in decades. Both osteocytes and adipocytes are endocrine cells regulating function of other organs. Osteocytes produce osteokines including sclerostin and FGF23, while adipocytes produce adipokines including adiponectin and leptin ^1,3^. Most importantly, both cell types are under transcriptional control of PPARG nuclear receptor ^6–8^.

Adiponectin represents a multifunctional adipokine considered to be expressed specifically in adipose tissue including marrow adipocytes ^9,10^. Adiponectin levels in circulation are substantial and they inversely correlate with obesity and insulin resistance. Adiponectin regulates fatty acids metabolism and insulin sensitivity in muscle, liver and adipose tissue, and hematopoiesis in the bone marrow. It also aids new bone formation during fracture healing ^11,12^. The progenitor cells of Adipoq-lineage have been identified in the bone marrow ^13–15^, and several reports have indicated that osteoblasts and osteocytes may express and produce adiponectin, however this finding has not been systematically pursued ^16–19^. During an investigation of the role of PPARG in regulation of osteocyte function, our group identified adiponectin as expressed in osteocytes under control of PPARG. This finding guided us to explore adipocyte-like phenotype of osteocyte and speculate on the functional meaning of such capabilities. The presented study concludes that PPARG is controlling adipocyte-like traits of osteocytes including adiponectin secretion at the levels suggesting contribution to the circulating adiponectin, and osteocyte capabilities to store and metabolize lipids which correlate with expression of perilipin 1 (PLIN1), fatty acids binding protein 4 (FABP4), hormone-specific lipase (HSL) and adipocyte-specific triglyceride lipase (ATGL). Physiologically, the adipocyte-like traits in osteocytes can be viewed as an adaptation to the isolated bone compartment in which they reside and the necessity to be self-relying on the accumulated fuel for survival and functional fitness.

## MATERIAL AND METHODS

### Generation of mice

The osteocyte-specific PPARγ-KO (OT^γKO^) mouse model was created by crossing PPARγ^fl/fl^ mice having loxP sites flanking the of exons 1 and 2 with Dmp1-Cre mice, as previously described ^7^. In all experiments, littermates with *Dmp*1^Cre^*Ppar* ^+/+^ genotype were used as control (Ctrl). The animals were maintained under 12hrs dark-light cycle with ad libitum access to water and chow, either regular (Teklad global 16% protein rodent diet; code:2916) or breeding (Teklad global 19% protein extruded rodent diet: code:2919). The breeding and experimental protocols (#107229 and #105923, respectively) conformed to the NIH National Research Council’s Guide for the Care and Use of Laboratory Animals and were reviewed and approved by the University of Toledo Health Science Campus Institutional Animal Care and Utilization Committee. The University of Toledo animal facility is operating as a pathogen-free, it is AAALAC approved facility and animal care and husbandry meet the requirements in the Guide for the Care and Use of Laboratory Animals.

Rosiglitazone supplemented diet experiment: twenty-two 3-month-old male mice (Ctrl and OT^γKO^) were randomly allocated to two groups and fed with either regular or rosiglitazone (Rosi) (Scripps Research Institute, Jupiter, FL) supplemented at the dose 25 mg/kg/day diet, as described previously^7^. After 8 weeks of treatment, animals were sacrificed, blood serum was harvested and osteocytes from cortical femora bone were isolated.

### Extraction of osteocyte-enriched fractions from femora bone and bone organ culture

Cell fraction enriched in osteocytes was isolated by sequential collagenase digestion of femora bone according to the previously described protocol^20^ ^7^. In brief, isolated femurs were cleaned and scraped to remove soft tissues and periosteum, respectively. Epiphyses were removed and bone marrow spun down. Bone diaphyses were sequentially digested in buffers supplemented with collagenase or EDTA^7^. Fraction number 6 representing osteocytes was lysed with TRI Reagent and total RNA and protein were extracted from the samples according to the manufacturer’s protocols (Sigma-Aldrich).

For organ cultures femoral bones underwent cleaning, removal of bone marrow and sequential collagenase digestion to remove periosteal and endosteal cells followed by bone fragmentation and seeding on the 30 mm dish in serum free αMEM media for 48 hrs. After collection, media were spun to remove cells and bone debris and were subjected to either the Adiponectin ELISA (cat #MRP300; R&D) or the Proteome Profiler Mouse XL Cytokine Array (cat #ARY028; R&D) analysis.

### Cell lines and culture conditions

MLO-Y4 cells edited with CRISPR/Cas9 for PPARγ deletion were developed using editing system designed by Synthego CRISPR Gene Knockout Kit v2 (Synthego Corporation, Redwood City, CA; Cat # SO6765399) and selected single cell-derived clones were verified for the editing efficiency using ICE (Interference of CRISPR Edits), an online bioinformatic software provided by Synthego. In all experiments, for the scientific rigor two separate clones with PPARγ KO (Y4^KO^) were used and compared to the control clone with mock CRISPR/Cas9 editing (Y4^C^). Cells were incubated at 37°C on collagen-coated plates, as previously described ^8^. Development and characterization of AD2 and OB6 cell lines representing marrow adipocytes and osteoblasts, respectively, were previously described^21^. MC3T3 cells were obtained from the American Tissue Culture Collection (ATCC). MLO-Y4 cells were obtained from Dr. Bonewald (Indiana University) and were cultured in α-MEM supplemented with 5% Fetal Bovine Serum (FBS), 5% Calf Serum, and 1% PS. All other cell lines were cultured in α-MEM supplemented with 10% FBS and 1% PS, at 37°C and the presence of 5% CO2. For rosiglitazone treatment, cells were grown to 70% confluency and treated with 10 µM rosiglitazone or DMSO as vehicle control, for 3 days following either RNA or protein isolation.

### BODIPY staining for lipids

For lipid droplet visualization 60,000 cells were seeded per well into 6-well plates containing 25 mm sterilized coverslips and cultured for 72 hrs with one media changed. To induce lipid droplet formation, cells were treated with 10 μM rosiglitazone (or DMSO vehicle control) for 48 hours prior to staining with BODIPY^TM^ 558/568 (Invitrogen) at 5 µM or 10 µM concentrations for 20 minutes at 37 °C. Cells were counterstained with 1 µg/ml DAPI for 2–3 minutes at room temperature to visualize nuclei. Coverslips were mounted onto glass slides using aqueous mounting medium and fluorescence imaging was performed using a confocal microscope and 63× oil-immersion objective.

### Fuel preference assay

Fuel preference of CRISPR/Cas9 *Ppar*γ edited cells (Y4^KO^) and mock-edited cells (Y4^C^) was measured using Seahorse Mito-Fuel Flex test (Agilent Technologies), in assay medium according to the manufacturer protocol. Cells were plated on assay micro-plate at the density of 20,000 cells per well. The assay uses three inhibitors: UK5099 (an inhibitor of the glucose oxidation pathway), BPTES (an inhibitor of the glutamine oxidation pathway), Etomoxir (an inhibitor of long chain fatty acid oxidation) provided in the Seahorse XF Mito fuel flex test kit (Agilent Technologies, Cat#103260-100). Fuel dependency for a specific pathway was measured after inhibiting this pathway. Fuel capacity of that pathway was measured after inhibiting two other pathways with specific inhibitors. Each measurement was performed on 8 replica wells. Software Wave version 2.6.1 was used for running the assay and obtaining data.

### Gene expression analysis using Quantitative Real-time RT-PCR

RNA (0.2 μg) was converted to cDNA using the Verso cDNA synthesis kit (Thermo Fisher Scientific, Waltham, MA). PCR amplification of the cDNA was performed by quantitative real-time PCR using TrueAmp SYBR Green qPCR SuperMix (Smart Bioscience, Maumee, OH) and processed with StepOne Plus System (Applied Biosystems, Carlsbad, CA). The thermocycling protocol consisted of 10 min at 95 °C, 40 cycles of 15 s at 95 °C, 30 s at 60°C, and 20 s at 72 °C, followed by melting curve step with temperature ranging from 60 to 95 °C to ensure product specificity. Relative gene expression was measured by the comparative CT method using 18S RNA levels for normalization. Primers were designed using Primer-BLAST (NCBI, Bethesda, MD) and are listed in Table 1.

**Table 1.**
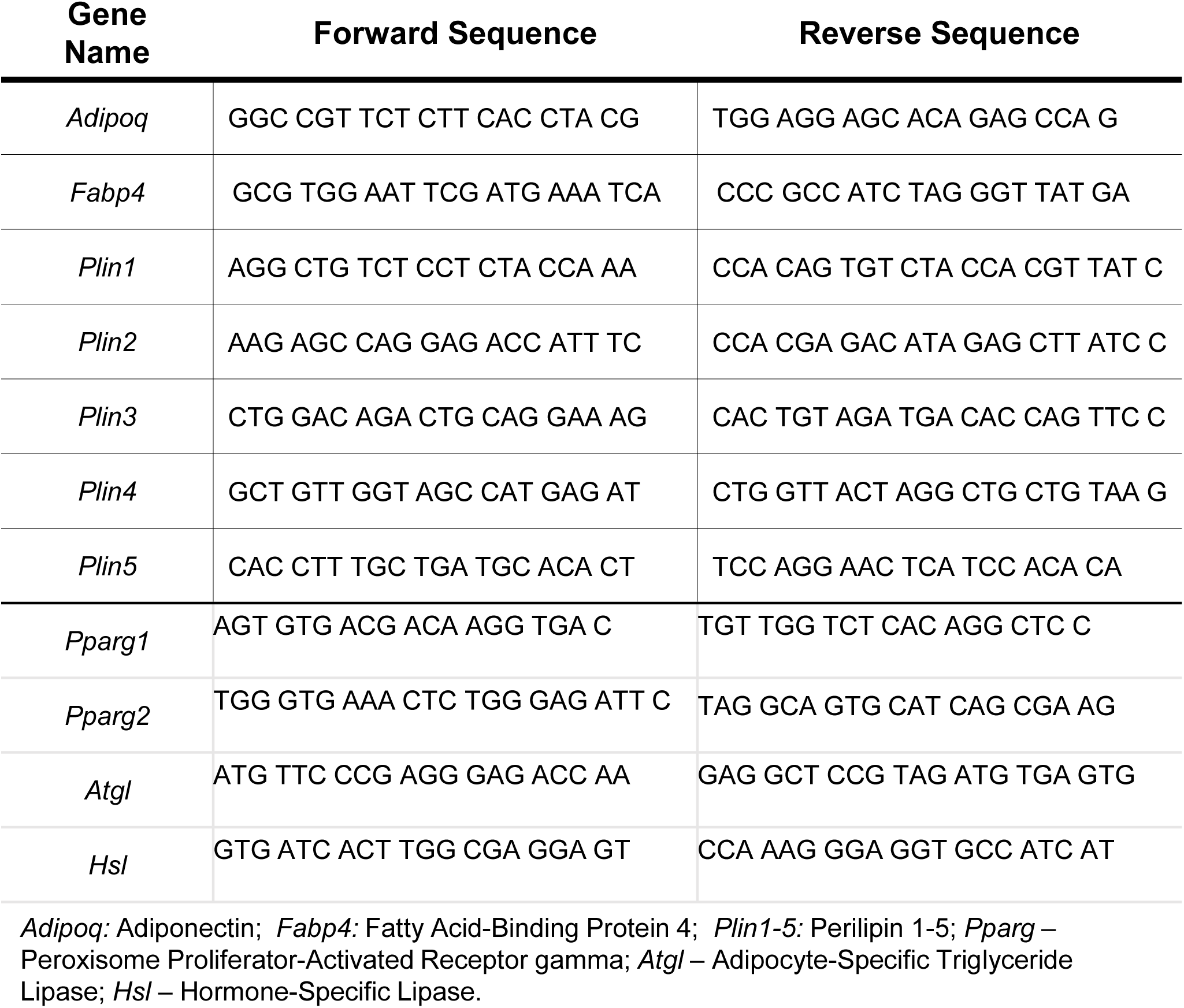

### Protein analysis using western blots

Protein fractions from MLO-Y4 osteocytes and AD2 adipocytes were isolated using lysis buffer containing 20nM Tris pH 7.5, 150mM NaCl, 1mM EDTA, 1% Triton (by weight), 2.5mM sodium pyrophosphate, 1mM beta-glycerophosphate, and protease and phosphatase inhibitors. 15 µg of lysate in Laemmli sample buffer was heated at 95°C for 5 minutes then loaded onto a 10% SDS-PAGE gel and ran at 115 volts for 1-1.5 hours. Proteins were transferred to a PVDF membrane which was blocked with 5% BSA for 1 hour. Incubation with primary antibody occurred overnight at 4°C in 1:1000 dilution, followed by secondary antibody at a 1:10,000 dilution for 1 hour. Membranes were washed and developed using the ECL method (Thermo Fisher Scientific, Waltham, MA 34577). Imaging was performed on a Syngene GBOX Chemi XX6 (Syngene, Frederick, MD 21704). Antibodies used for Western blots: PPARG Rb monoclonal (cat#81B8; Cell Signaling Technologies), β-Actin Ms monoclonal (A1978; Sigma), Anti-Rabbit IgG HRP-linked (cat#7074; Cell Signaling Technologies), and Anti-Mouse IgG HRP-linked (cat#sc-2005; Santa-Cruz Biotechnology).

### Statistical Analysis

Data are represented as means ± SD and were analyzed by using GraphPad Prism 9.1. A parametric, unpaired, two-tailed Student’s t-test was used to compare the means of datasets with two groups. One-way ANOVA followed by Tukey’s multiple comparison tests was used to compare means of datasets with two or more groups. Pearson correlation coefficients were computed with 95% confidence interval and two-tailed p value. Statistically significant p values are indicated by asterisks as follows: * p<0.05, ** p<0.01, *** p<0.001, and ****p<0.0001.

## RESULTS

### Osteocytes produce and secrete adiponectin under control of PPARG

Adiponectin is a cytokine considered to be expressed predominantly in adipocytes. Our analysis of the bulk transcriptomics of osteocyte fraction isolated from mice femora cortical bone suggested that adiponectin transcript (*Adipoq*) is also expressed in osteocytes in a relatively high level ^22^. To follow this finding, we examined levels of adiponectin (ADIPOQ) protein expression in femora cortical bone of 8 mo old C57BL/6 female and male mice and compared it to the levels of expression in the interscapular brown (BAT) and gonadal/epidydimal white (WAT) adipose tissues isolated from the same mice. In addition, ADIPOQ levels were analyzed in the bone marrow, which at this age is substantially infiltrated with marrow adipocytes. After normalization to gram unit of analyzed tissue, ADIPOQ protein production in cortical bone, which after isolation had been processed for osteocyte enrichment, as described in Material and Methods, was at similar levels as in BAT and much higher than in WAT and bone marrow (Fig. 1A). As shown in ex vivo bone organ culture, ADIPOQ protein is not only produced, but it is also secreted from cortical bone with retained sex-specific ratio (Fig. 1B). However, when calculated per mass of two femora bones and compared to production in interscapular BAT, the amount of ADIPOQ protein produced by femora is of a magnitude lower than in BAT (Fig. 1C) suggesting rather modest contribution to the overall levels of circulating ADIPOQ.

**Figure 1.**
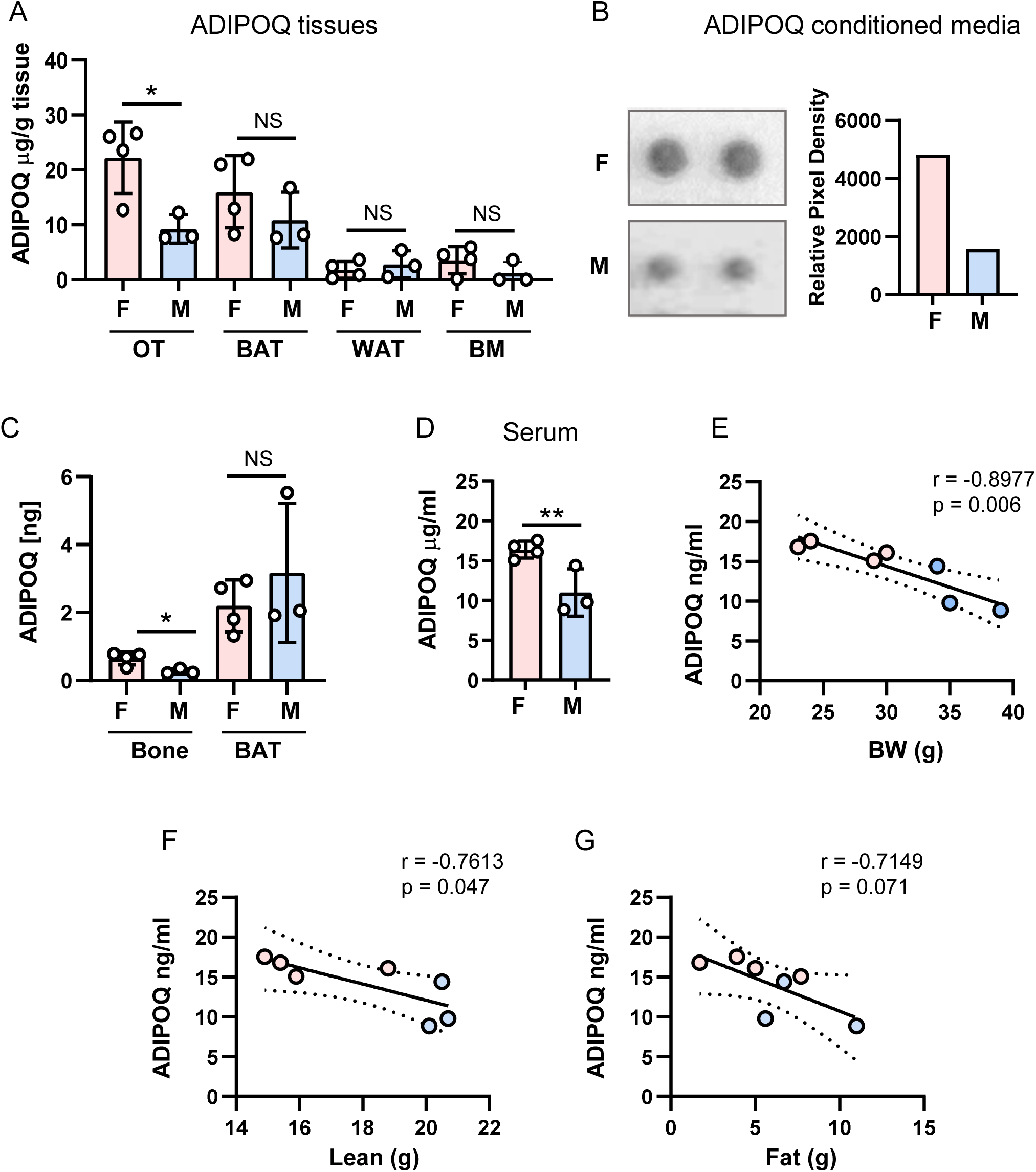
Adiponectin (ADIPOQ) production in different tissues including bone and its levels in circulation, in 8 mo old female (F - pink) and male (M - blue) C57BL/6 mice. A. ADIPOQ production measured by ELISA diagnostic kit and normalized per gram of that tissue. Osteocytes (OT) were isolated from femora cortical bone, as described in Material and Methods. WAT, white adipose tissue, represents gonadal and epididymal fat depots, BAT represents interscapular brown adipose tissue, and BM represents protein lysates of bone marrow isolated from the analyzed femora bone. B. ADIPOQ detection in conditioned media collected from organ culture of enriched for osteocytes cortical bone derived from 5 mo old female and male using Proteome Profiler Mouse XL Cytokine Array. Graph represents a relative pixel density of replicate dots corresponding to ADIPOQ on the Array. C. Calculated ADIPOQ production in two femora compared to entire interscapular BAT of the same 8 mo old F and M. D. Serum levels of ADIPOQ in 8 mo old F and M. Statistical significance was calculated using two-tailed Student T-test. * p < 0.05; ** p < 0.01; NS – not significant. E – G. Pearson correlation analysis of relationship between ADIPOQ serum levels and an animal body weight (E), a weight of lean tissue (F), and a weight of fat tissue (G) measured by NMR. Pearson correlation coefficient (r) and significance (p) are listed on graph in panels E – G.

An analysis of sera isolated from the same mice as presented in Fig. 1A-C showed higher levels of circulating ADIPOQ in females (Fig. 1D). Interestingly, the same sexual divergence is also seen in osteocytes. The *Adipoq* expression, and ADIPOQ production and secretion from female cortical bone exceeded by 2-fold the levels in male cortical bone, while these sex-specific differences were not observed in analyzed fat depots including BAT, WAT and BM of the same animals (Fig. 1A-C). Moreover, ADIPOQ levels correlated negatively with an animal body weight (Fig. 1E) and a weight of lean tissue (Fig. 1F), but not with a weight of fat tissue measured by NMR (Fig. 1G). On the note, the females body weight and lean mass were substantially lower than males (Fig. 1F and E, blue vs. pink symbols) emphasizing sexual divergence in ADIPOQ production.

Similarly to adipocytes, ADIPOQ RNA and protein expression in osteocytes are under PPARG control (Fig. 2). An analysis of *in vivo* osteocyte fraction isolated from femora cortical bone of mice with osteocyte-specific deletion of PPARG ((Dmp1^Cre^PPARγ^flfl^ or γOT^KO^) and compared to Ctrl mice (Dmp1^Cre^PPARγ^+/+^) confirmed that *Adipoq* is under positive transcriptional control of PPARG (Fig. 2A). This correlated with a significantly lower expression of ADIPOQ protein in cortical bone (Fig. 2B) and it was correlated with 10% lower levels of ADIPOQ protein in circulation of γOT^KO^ mice (Fig. 2C). To confirm that PPARG nuclear receptor positively controls *Adipoq* expression, Ctrl and γOT^KO^ mice were treated with Rosiglitazone (Rosi), an anti-diabetic drug with full agonist activity for PPARG ^9^. In vivo treatment with Rosi increased by 4-fold *Adipoq* expression in femoral bone osteocytes of Ctrl mice, while this effect was abolished in osteocytes present in cortical bone of γOT^KO^ mice (Fig.2D). Measurement of ADIPOQ levels in circulation reflected this observation. Although in both γOT^KO^ and Ctrl mice treated with Rosi ADIPOQ levels increase, which accounts for Rosi effects mainly on adipose tissues, the increase in sera of γOT^KO^ mice did not reach the same levels as in Ctrl mice (Fig. 2E) and were lower by approximately 10%. These suggest that bone contributes to the circulating ADIPOQ levels that this contribution is under control of PPARG which activity can be pharmacologically modulated in skeletal tissue.

**Figure 2.**
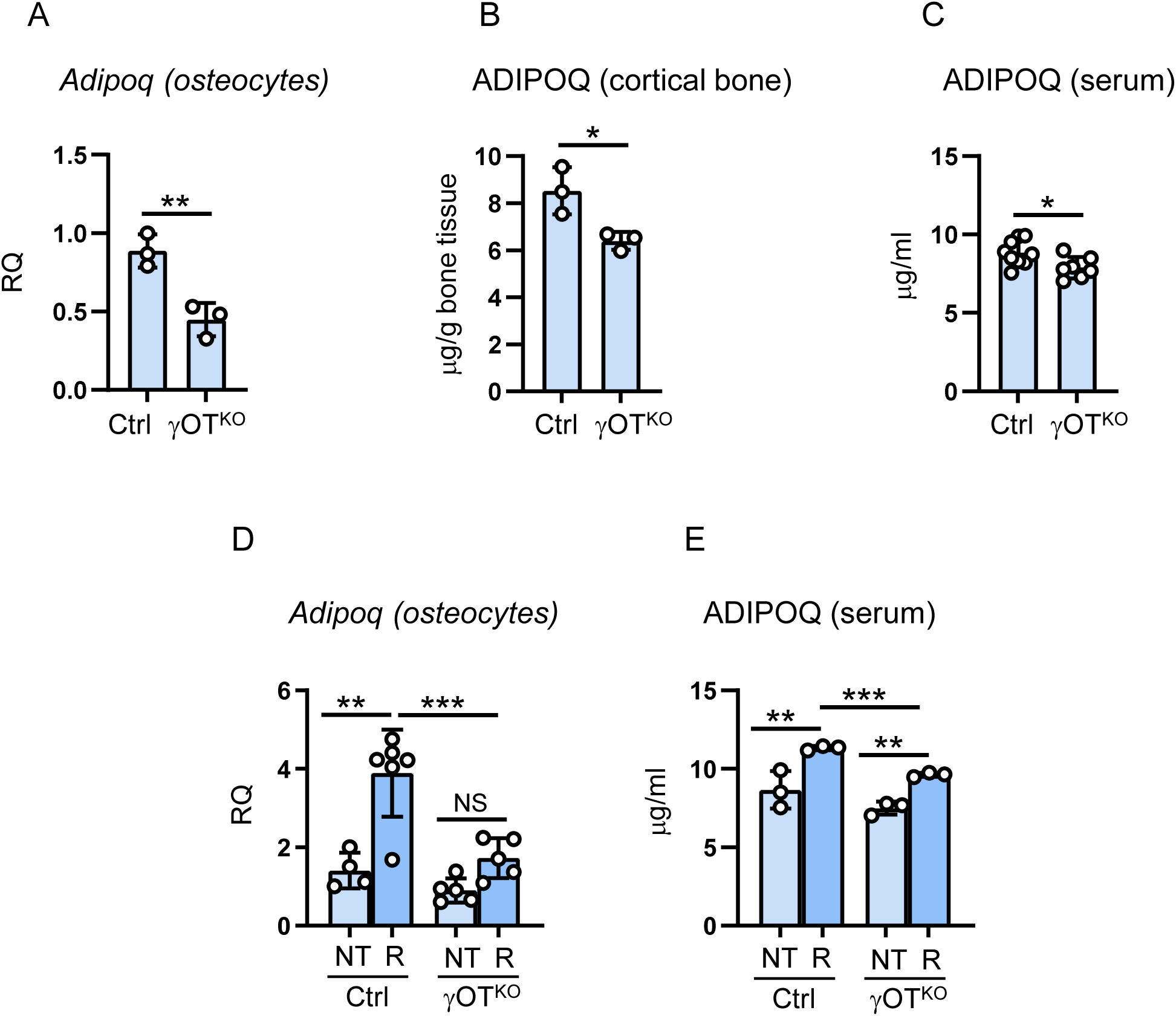
*Adipoq* mRNA expression and ADIPOQ protein levels in in vivo osteocytes isolated from cortical femoral bone and serum of 6 mo old Ctrl and γOT^KO^ male mice with osteocyte-specific deletion of PPARG. A. Levels of *Adipoq* in *in vivo* osteocytes. B. ADIPOQ protein levels in osteocyte-enriched cortical bone homogenates. C. ADIPOQ serum levels. D. *Adipoq* expression in in vivo osteocytes after 8-week treatment with PPARG agonist rosiglitazone, as described in Material and Methods (NT – not treated, R – rosiglitazone treated). E. ADIPOQ serum levels in animals as in D. Statistical significance was calculated using two-tailed Student T-test. * p < 0.05; ** p < 0.01; *** p < 0.001.

In overall, presented data indicate that osteocytes express and secrete adiponectin, that this expression is at the level of expression in brown adipocytes, that in osteocytes levels of ADIPOQ expression follows sex-specific pattern seen in the circulation, that ADIPOQ expression is under control of PPARG, and that osteocytes secrete ADIPOQ at the levels which can contributes to the pool of circulating adiponectin.

### Osteocytes accumulate lipids under PPARG control

Two major features of adipocytes are their endocrine activity signified by adipokines production and storage of lipids in the form of triglycerides in specialized vacuoles. Lipid vacuoles metabolism is controlled by number of proteins including perilipins. These proteins, with a prominent role of Perilipin 1 in adipocytes and Perilipin 2 in osteoblasts, are responsible for storage and lipolysis of triglycerides ^23^. Thus, next step in our examination of osteocyte adipocytic characteristics consisted of analyses of their ability to accumulate lipids as a function of PPARG and whether this was associated with transcriptional changes in the expression of perilipin proteins.

Using BODIPY^TM^ 558/568 fluorescent staining of neutral lipids, we found that MLO-Y4 osteocytes are naturally capable of accumulating and storing lipids in vacuoles located in the cytoplasm (Fig. 3A). Most importantly, this ability depends on the presence of PPARG protein because MLO-Y4 cells edited for PPARγ with CRISPR/Cas9 are unable to form lipid vacuoles, despite the lipids being present in the cytoplasm (Fig. 3A).

**Figure 3.**
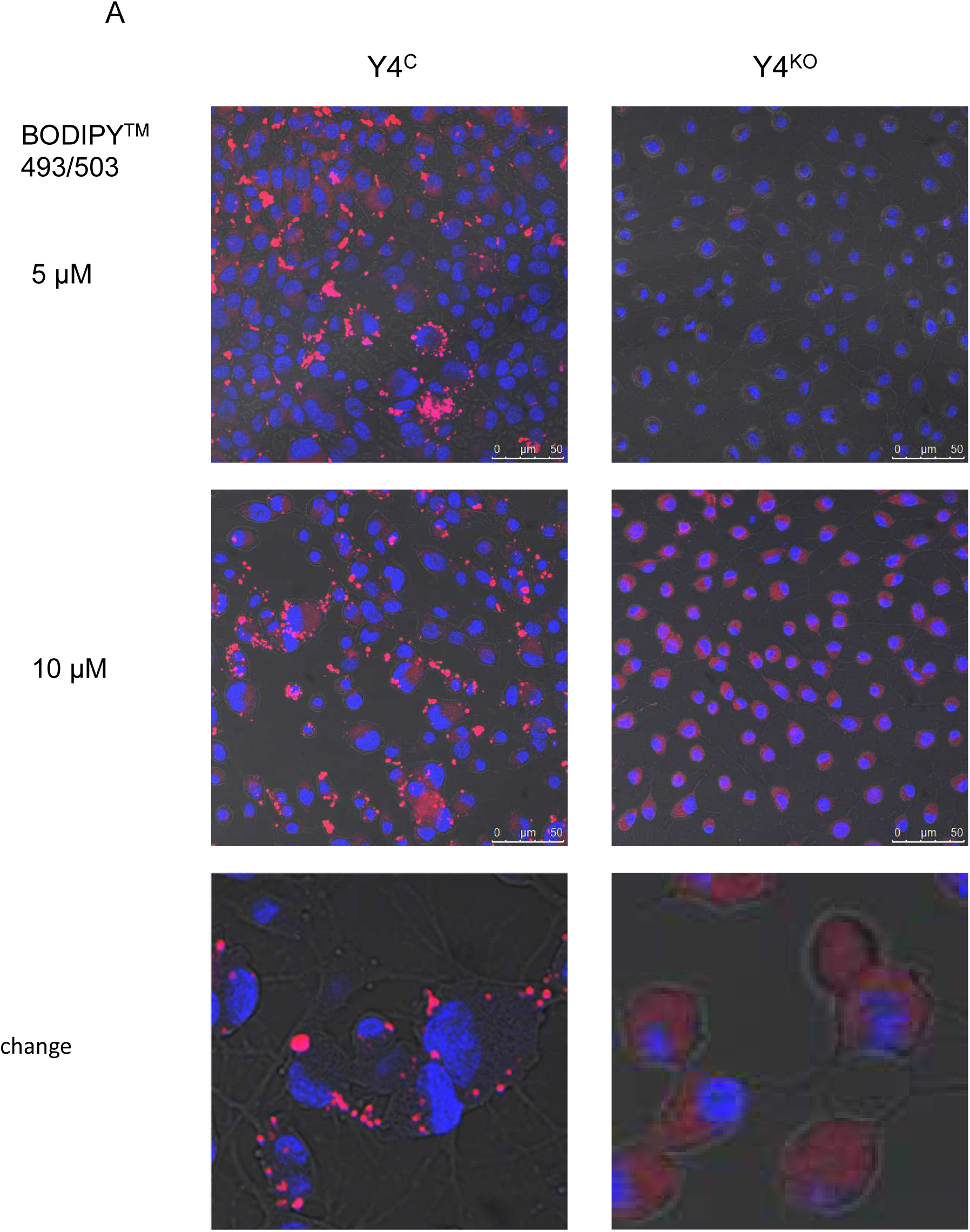
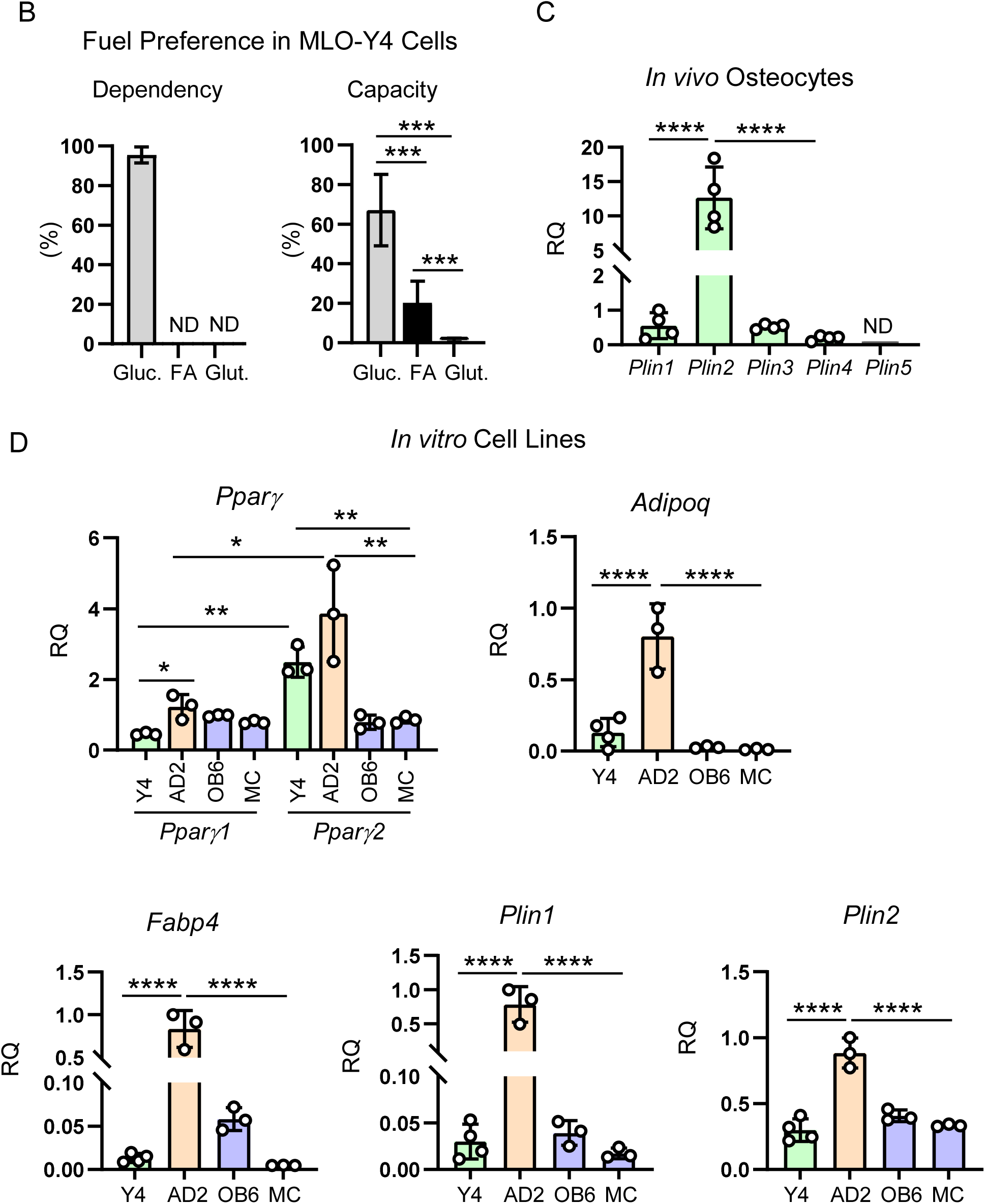
Osteocytes accumulate and use lipids and express adipocyte-specific gene markers under control of PPARG. A. Visualization of lipid droplets in MLO-Y4 cells mock-or PPARG-edited with CRISPR/Cas9 (Y4^C^ and Y4^KO^, respectively). Imaging using confocal microscopy, magnification 63×, lipids stained with BODIPY^TM^ 493/503, as described in Material and Methods. Scale bars indicate 50 µm. B. Fuel preference (dependency and capacity) of MLO-Y4 cells measured with Seahorse FuelFlex assay as described in Material and Methods. Gluc. – glucose, FA – fatty acids, Glut. – glutamine. C. Relative expression of Perilipin family transcripts in osteocytes isolated from 3 mo old C57BL/6 male mice. Levels of expression *Plin 2-5* were normalized to *Plin1*. D. Relative levels of adipocyte-specific gene markers expression in MLO-Y4 osteocytes (Y4), OB6 and MC3T3 (MC) osteoblasts, and AD2 marrow adipocytes to which all values were normalized. Statistical significance was analyzed with One-way Anova with multiple comparison. * p < 0.05; ** p < 0.01; *** p < 0.001; **** p < 0.0001. ND – not detected.

There is an ongoing discussion on osteocyte bioenergetics, specifically the use of fatty acids as a fuel source. Using the Agilent Seahorse XF Mito Fuel Flex Test we tested a dependency and capacity of MLO-Y4 cells to oxidize three critical mitochondrial fuels: glucose, fatty acids, and glutamine. This test determines the rate of oxidation in basal state of each fuel by measuring mitochondrial respiration (OCR) in the presence of selective inhibitors such as UK5099, Etomoxir and BPTES which block oxidation of glucose, long chain-fatty acids, and glutamine, respectively. The assay measures both fuel dependency which informs of cells’ reliance on a certain fuel to maintain basal respiration, and fuel capacity which informs of cells’ ability to meet metabolic demand from one fuel when other fuel pathways are inhibited. As shown in Fig. 3B, in basal conditions MLO-Y4 cells rely almost entirely on glucose, with negligible contribution of fatty acids and glutamine as energy source. However, when glucose and glutamine use is blocked by specific inhibitors, osteocytes are capable to use fatty acids for covering up to 20% of their energy requirement (Fig. 3C).

An analysis of the RNA expression of Perilipin family members, showed that with the exception of Perilipin 5 (*Pln5*), which has been previously identified as specifically expressed in highly oxidative tissues including brown adipose tissue (BAT) and muscle ^23^, transcripts for *Pln1-4* are present in *in vivo* osteocytes (Fig. 3C). However, unlike adipocytes which highly express *Pln1*^23^, in osteocytes *Pln2* is the most highly expressed. It has been previously shown that *Pln2* is specifically expressed in osteoblast where it regulates lipid metabolism ^24^. Thus, high expression of *Pln2* in osteocytes is consistent with them being the descendants of osteoblasts.

Further, we compared the level of expression of adipocyte-specific gene markers including their transcriptional regulator *Ppar*γ, in established cell lines representing osteocytes (MLO-Y4), marrow adipocytes (AD2), and osteoblasts (OB6 and MC3T3). PPARG protein consists of two isoforms, G1 and G2, which are products of the same gene transcript originating from an alternative promoter and being alternatively spliced. While PPARG1 isoform is ubiquitously expressed in many cell types, PPARG2 is specifically expressed in cells able to store lipids such as adipocytes or “foam” macrophages ^25^. As shown in Fig. 3D, the expression of *Ppar*γ*2* is significantly higher in MLO-Y4 and AD2 cells than expression of *Ppar*γ*1*, which is consistent with MLO-Y4 cells adipocyte-like phenotype. Notably, the expression of *Ppar*γ*1* and *Ppar*γ*2* transcripts is on a similar and relatively low levels in osteoblastic OB6 and MC3T3 cells.

Subsequent comparison showed that AD2 cells have the highest expression levels of tested adipocyte-specific gene markers including *Adipoq*, *Fabp4*, *Pln1* and surprisingly *Pln2* (Fig. 3D), which might reflect their origin from a common mesenchymal progenitor with osteoblasts. In this analysis, the basal expression of adipocyte markers, with exception to *Adipoq*, was on the same low levels in MLO-Y4, OB6, and MC3T3 cells, as compared to AD2 cells in which these levels were from 4-to 20-fold higher.

### Osteocytes are much like adipocytes, but not like osteoblasts, in their response to rosiglitazone-inducing PPARG pro-adipocytic activity

PPARG pro-adipocytic activity is induced efficiently with Rosi which acts as a full agonist for this nuclear receptor. Similarly to the autoregulatory activity in extramedullary adipocytes, PPARG protein levels increase upon Rosi treatment in MLO-Y4 osteocytes with the same magnitude and almost to the same levels as in untreated AD2 marrow adipocytes (Fig. 4A). Consequently, treatment with Rosi of MLO-Y4 cells increased number of lipid droplets although not as robust as in AD2 cells (Fig. 4B and C). In contrast, treatment with Rosi of MLO-Y4 cells deficient in PPARG and treatment of OB6 cells did not invoke such response.

**Figure 4.**
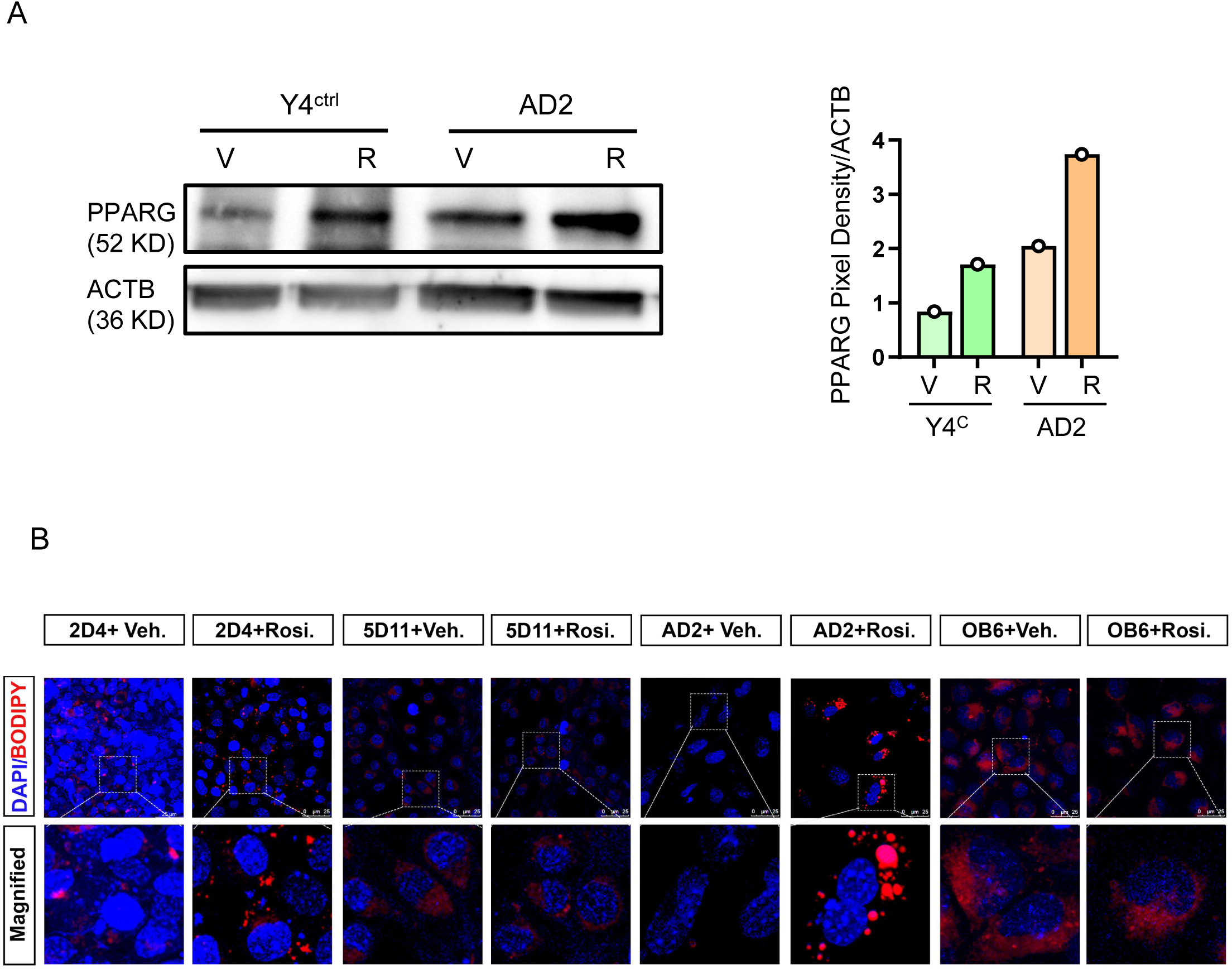
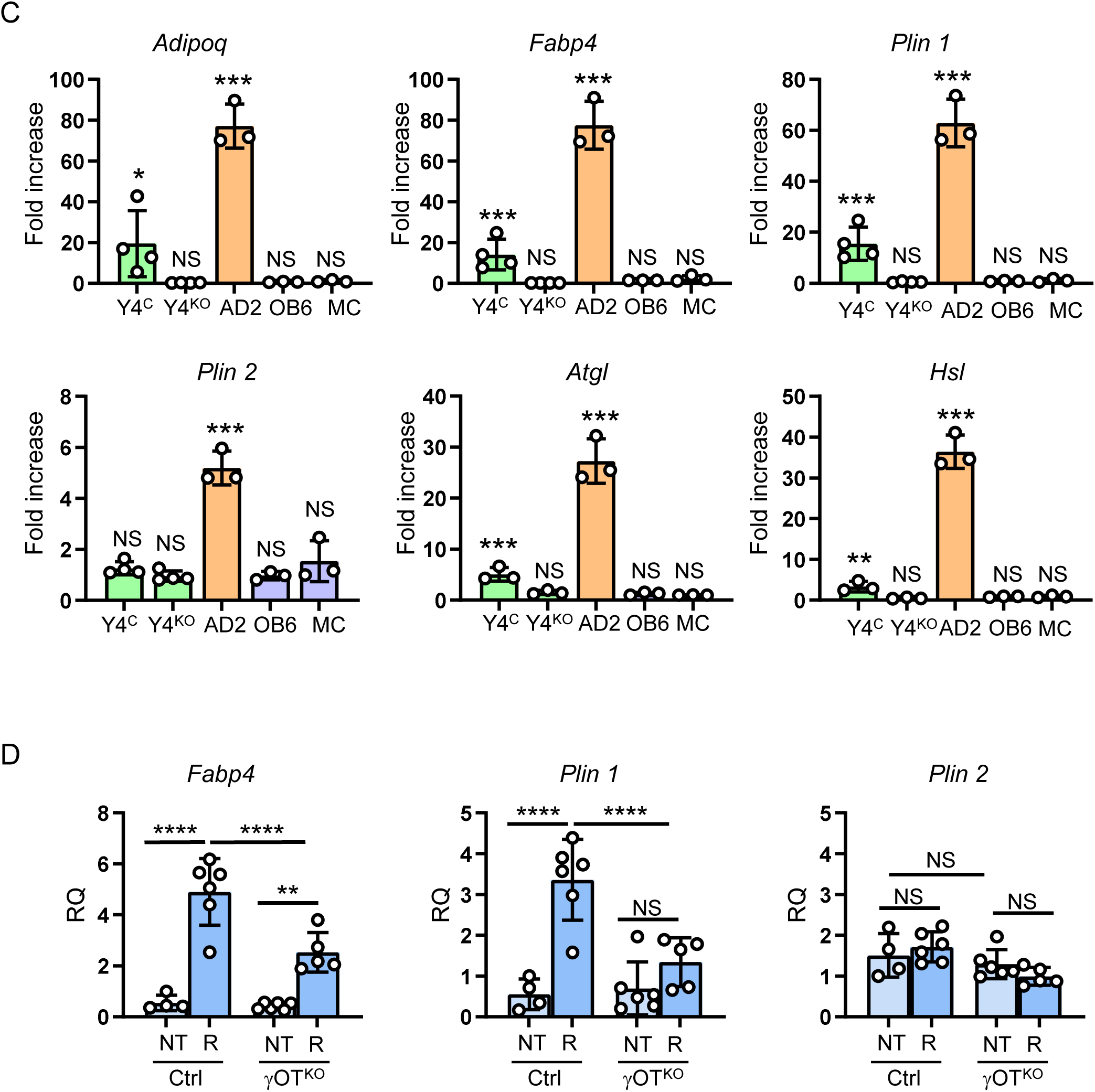
Osteocytes, but not osteoblasts, respond to pro-adipocytic activity of rosiglitazone as a function of PPARG protein. A. Western blot of PPARG levels in MLO-Y4 and AD2 cells in response to rosiglitazone treatment. Graph on the right represents relative pixel density of bands representing PPARG after normalization to levels of β-actin protein (ACTB). B. Fluorescent staining with BODIPY^TM^ 493/503 for lipids in MLO-Y4 osteocytes (Y4^C^ and Y4^KO^ cells), AD2 marrow adipocytes, and OB6 osteoblasts after treatment with 10 µM rosiglitazone or vehicle DMSO for 3 days. C. Fold of induction of adipocyte-specific gene markers in cells treated with 10 µM rosiglitazone, as normalized to DMSO-treated cells serving as control. Statistical significance was calculated using two-tailed Student T-test. * p < 0.05; *** p < 0.001. D. Expression of *Fabp 4*, *Plin 1* and *Plin 2* in in vivo osteocytes isolated from femora bone of γOT^KO^ and Ctrl mice receiving diet supplemented with rosiglitazone for 8 weeks, as described in Material and Methods. (NT – not treated, R – rosiglitazone treated). Statistical significance was analyzed with One-way Anova with multiple comparison. ** p < 0.01; *** p < 0.001; **** p < 0.0001; NS – not significant.

The levels of induction of adipocyte-specific gene markers of MLO-Y4 cells in response to Rosi treatment was consistent with the pro-adipocytic response of these cells. In contrast to osteoblastic OB6 and MC3T3 cells, MLO-Y4 cells responded with increased expression of *Adipoq*, *Fabp4*, *Pln 1*, *Atgl*, and *Hsl* transcripts, albeit with a lower magnitude than in AD2 cells (Fig. 4D). The expression of these transcripts in MLO-Y4 cells was at the range of 4-to 20-fold compared to 20-to 80-fold in AD2 cells, indicating that osteocytes are in the spectrum of adipocyte phenotype. Notably, osteocyte response to Rosi treatment was PPARG-dependent since γY4^KO^ cells did not response to treatment (Fig. 4D).

### Systemic administration of rosiglitazone to animals induces adipocyte-like phenotype in cortical osteocytes as a function of PPARG activity

To prove physiological relevance, the pro-adipocytic activity of PPARG in osteocytes was tested *in vivo*. RNA transcripts isolated from cortical osteocytes of male mice, with either intact PPARG (Ctrl) or deleted specifically in osteocytes (γOT^KO^), fed for 8 weeks a diet supplemented with Rosi, were tested for expression of adipocyte-specific gene markers. In addition to increased expression of *Adipoq*, previously shown in Fig. 2D, the expression of *Fabp4* and *Plin1* was markedly increased in Ctrl animals receiving Rosi (Fig. 4E). This response was either blunted or of substantially lower magnitude in osteocytes derived from γOT^KO^ mice. Upon Rosi administration, no significant differences in expression of *Plin2* was noted (Fig. 4E), as well as for *Plin 3* and *4*, and transcript for *Plin5* was still not detected (data not shown), suggesting that in osteocytes these transcripts are not under control of PPARG pro-adipocytic activity. Taken together, presented data suggest that osteocytes are “predisposed” to invoke the pro-adipocytic activity *via* PPARG nuclear receptor.

## DISCUSSION

Presented studies show that osteocyte phenotype partially consists of adipocyte-like characteristics that are under control of PPARG nuclear receptor, manifested in the unique osteocyte-specific manner. These include production and secretion of adiponectin and marked intracellular lipid accumulation and metabolism, while retaining their osteocytic phenotype. These findings provide new insight into osteocytes’ functional link to the network controlling systemic energy metabolism.

Our study shows that adiponectin, the adipocyte flagship cytokine, is produced and is secreted by osteocytes in a quantity significant for local function and, albeit modest, contribution to the circulating levels of this cytokine. Our data collected from the bone of the same animal shows that the major source of adiponectin produced in murine adult bone are osteocytes, while production from marrow adipocytes is negligible. Moreover, adiponectin production in osteocytes is sex dependent, higher in females than in males, which is in contrasts with adiponectin production in WAT and BAT of the same mouse. The correlation analysis confirmed that sexual differences in circulating levels of adiponectin relate to lean not to fat content of body composition. It is well documented in humans that females have higher levels of adiponectin in circulation than men with testosterone being considered the main negative determinant of adiponectin production^26,27^. If adiponectin expression in osteocytes is modulated by sex hormones, this may provide an additional target to modify osteocyte function controlling bone homeostasis and bone repair, as discussed below.

Adiponectin orchestrates a broad spectrum of biological and biochemical processes, including lipid metabolism, energy homeostasis, inflammatory regulation, and insulin sensitivity. Its levels in circulation are negatively associated with obesity and insulin resistance ^9^. It has been demonstrated *in vitro* and in animal studies that adiponectin has a positive effect on bone formation and negative effect on bone resorption ^16,28–31^. When applied at the site of injury, adiponectin facilitates bone regeneration and fracture healing ^11,12^. These findings underscore the beneficial effects of local, paracrine activity of adiponectin. However, adiponectin’s role in bone is still intriguing and complex ^32^. Both of its receptors, AdipoR1 and AdipoR2, which respond to either globular or high molecular weight of adiponectin protein, are expressed in primary osteoblasts, marrow adipocytes and marrow macrophages^33^. The adiponectin mimetic GTDF (flavonol 6-C-β-D-glucopyranosyl (2S,3S)-(+)-5,7,3′,4′-tetrahydroxydihydroflavonol) with a functional bias toward AdipoR1 normalizes osteopenia in db/db mice lacking AdipoR1 expression in skeletal muscle but not in bone cells, independently of systemic glycemic correction of type 2 diabetes phenotype ^34,35^. Mechanistically, this skeletal protection is associated with mitigation of gluco-and lipotoxicity-induced osteoblast apoptosis linking adiponectin signaling to skeletal integrity and bone cell survival ^36^. Thus, it is of high interest to define the role of adiponectin not only in the context of osteocyte endocrine activity, but also for a possibility that in autocrine manner it regulates osteocyte function and survival.

A display of markers of fatty acid metabolism by osteocytes in basal as well as stimulated conditions confirms their utilization for energy production. We have previously shown that under control of PPARA nuclear receptor lipid metabolism in osteocytes is of such magnitude that its impairment alters a balance of systemic energy metabolism and leads to development of obesity ^37^. Similar effect is observed with the inhibition of fatty acids β-oxidation due to CPT2 deletion in these cells ^38^ indicating that osteocytes are important entity in systemic lipid utilization.

Here, we showed that osteocytes metabolize lipids under control of PPARG in a manner similar to adipocytes but simultaneously specific to osteocytes. In these cells, basal expression of *Plin1* and *Fabp4* transcripts is relatively low and comparable to osteoblasts, which is in contrast with lineage committed adipocytes where they are highly expressed. However, upon induction of PPARG pro-adipocytic activity the expression of these transcripts in osteocytes is induced substantially and correlates with increased lipids accumulation in the form of lipid droplets. In these conditions, osteocytes also invoke an expression of lipolytic enzymes, ATGL and HSL, suggesting that accumulated lipids are used to support their metabolism. Notably, osteoblasts do not respond to pro-adipocytic treatment, although they express *Ppar*γ*1* and *Ppar*γ*2* transcripts. However, there is one distinction which may account for different responses of these two types of cells. In contrast to osteoblasts, the osteocytes express significantly higher levels of transcript coding for PPARG2 isoform, which we previously characterized as essential for marrow mesenchymal cells commitment toward adipocytic lineage ^21^. This suggests that during their transformation from osteoblasts, the osteocytes undergo reprogramming and acquire adipocyte-like features.

It is not a unique attribute for cells of mesenchymal origin to accumulate lipids, in both normal and pathological conditions. Lipid droplets accumulation in healthy muscle is associated with their fitness and response to exercise ^39^, while in diabetes it is a marker of impaired glucose utilization in response to insulin ^40^. However, what distinguishes muscle cells from osteocytes is that in muscle lipid droplets accumulation is under control of PLIN2 and their metabolism is under control of PLIN5 and PPARA nuclear receptor^41^, while in osteocytes and similarly to adipocytes, lipid accumulation is under control of PLIN1 and PPARG nuclear receptor. PLIN1 is primarily present in mature lipid-storing adipocytes and is located on the surface of lipid droplets. Besides acting as a scaffold for droplets, PLIN1 regulates lipolysis by facilitating ATGL activity ^42^.

Our data showing that although dependent on glucose, osteocytes have capacity to utilize fatty acids as a fuel source, supports recent notion on their metabolic flexibility ^43^. It has been demonstrated in vitro that differentiation of IDG-SW3 osteocytes is constrained by glucose dependency for anaerobic glycolysis to produce energy. In contrast, terminally differentiated osteocytes exhibit pronounced metabolic adaptability including compensation for the absence of glucose with fatty acid metabolism. Interestingly, this is accompanied by reduced expressions of *Sost* and *Fgf23*, suggesting that fuel use is functionally integrated into osteocyte biology. Moreover, mechanical loading enhances fatty acid β-oxidation pathways, indicating that osteocytes dynamically reprogram their metabolism in response to environmental and mechanical cues ^43^. Thus, our finding that osteocytes possess a lipid storage capacity and that their metabolism is reminiscent of adipocytes, supports a fundamental hallmark of metabolic flexibility in these long-lived, embedded cells.

These together provide a new understanding of osteocyte biology indicating simultaneous maintenance of osteoblast-like features as progeny of this lineage and acquisition of distinctive features of adipocytes, probably as an adaptation to the unique localization and a need for the maintenance of their fitness, functional adaptability and survival. Better understanding of the crossroads of osteocyte and adipocyte function may provide bases for development of therapeutic means to target common molecular pathways to manage metabolic disorders and skeletal diseases, simultaneously.

## AUTHORS’ CONTRIBUTION

MPK, EC, BLC – conceptualization, data curation, writing, reviewing;

MPK, EC, JL, SB, PJC - investigation, formal analysis, methodology, visualization.

## ACKNOWLEDGMENTS

This study was supported to BLC by the National Institute on Aging grant number R01AG071332.

